# The role of hybridization during ecological divergence of southwestern white pine (*Pinus strobiformis*) and limber pine (*P. flexilis*)

**DOI:** 10.1101/185728

**Authors:** Mitra Menon, Justin C. Bagley, Christopher Friedline, Amy V. Whipple, Anna W. Schoettle, Alejandro leal-Saenz, Christian Wehenkel, Francisco Molina-Freaner, Lluvia Flores-Renteria, M. Socorro Gonzalez-Elizondo, Richard A. Sniezko, Samuel A. Cushman, Kristen M. Waring, Andrew J. Eckert

**Author notes:** Authors for correspondence: Mitra Menon, 1000 West Cary Street, Integrative Life Sciences, Virginia Commonwealth University, Richmond, Virginia 23284 USA,; Andrew J. Eckert, 1000 West Cary Street, Life Science Bldg. - Department of Biology Rm 126, Virginia Commonwealth University, Richmond, Virginia 23284 USA,.

## Abstract

Interactions between extrinsic factors, such as disruptive selection, and intrinsic factors, such as genetic incompatibilities among loci, can contribute to the maintenance of species boundaries. The relative roles of these factors in the establishment of reproductive isolation can be examined using species pairs characterized by gene flow throughout their divergence history. We investigated the process of speciation and the maintenance of species boundaries between *Pinus strobiformis* and *P.flexilis*. Utilizing ecological niche modeling, demographic modeling, and genomic cline analyses, we illustrated a history of divergence with continuous gene flow between these species. We found an abundance of advanced generation hybrids and a lack of loci exhibiting large allele frequency differences across the hybrid zone. Additionally, we found evidence for climate-associated variation in the hybrid index and niche divergence between parental species and the hybrid zone. Our results are consistent with extrinsic factors, such as climate, being an important isolating mechanism for these species. A buildup of intrinsic incompatibilities and of co-adapted gene complexes is also apparent in our results, although these appear to be in the earliest stages of development. This supports previous work in coniferous species demonstrating the importance of extrinsic factors in creating and enforcing species boundaries. Overall, we lend support to the hypothesis that varying strengths and directions of selection pressures across the long lifespans of conifers, in combination with their life history strategies, delay the evolution of strong intrinsic incompatibilities.

## Introduction

Speciation often occurs along a continuum of divergence such that evolutionary processes leading to species formation initially involve unrestricted gene flow followed by the evolution of reproductive isolation between lineages (Kane *et al.* 2009; Nosil & Feder 2012; Roesti *et al.* 2012). Hence, understanding how and when barriers to gene flow arise and are maintained along this continuum is a fundamental goal of evolutionary biology (Losos *et al.* 2013). Under a model of ecological speciation (Schluter & Conte 2009), initiation of divergence among populations occurs through disruptive selection leading to the formation of ecotypes. This process results in shifts of allele frequencies correlated with environmental differences between habitats specific to each ecotype. The subsequent transition from ecotypes to reproductively isolated species occurs through the build-up of associations between several loci independently experiencing disruptive selection, and the action of selection to maintain these co-adapted divergent gene complexes (Flaxman *et al.* 2014).

Several studies of speciation have used hybrid zones as windows into the process of divergence among species (reviewed by Petit & Excoffier 2009). Studies conducted across the entire geographical range of hybridizing species have helped reveal not only the demographic context of speciation, but also the relative importance of intrinsic and extrinsic processes (Schield *et al.* 2017; Ryan *et al.* 2017). Specifically, the maintenance of species boundaries has been shown to occur through models of tension zones (intrinsic incompatibilities *sensu* Barton & Hewitt 1985; Via *et al.* 2000; Barton 2001; Rundle 2002) and bounded hybrid superiority (extrinsic incompatibilities *sensu* Moore 1977; Milne *et al.* 2003; Hamilton *et al.* 2013).The former facilitates divergence through a buildup of genetic incompatibilities among loci causing environmentally independent reduction in fitness of the hybrids, whereas the latter involves increased hybrid fitness only in a novel environment to which the divergent parental allelic combinations confer a putative advantage. These two models are often coupled, such that genomic regions involved in intrinsic incompatibility coincide with loci exhibiting ecological gradients in allele frequency (Bierne *et al.* 2011; Cushman & Landguth 2016), ensuring the maintenance of species barriers under the homogenizing effect of gene flow (Kulmuni & Westram 2017). Thus, the interaction between intrinsic and extrinsic barriers to gene flow generates a genomic mosaic of introgression and differentiation that depends in part upon the demographic context and life history traits of the diverging lineages.

The recent influx of genomic data from non-model species has facilitated studies of ecological speciation across varying spatial and temporal scales (Lexer *et al.* 2010; Andrew & Rieseberg 2013; de Lafontaine *et al.* 2015; Lackey & Boughman 2016; Marques *et al.* 2017). Studies using genome scans often lend support to the genic view of speciation (Wu 2001), which predicts that a handful of genomic regions experiencing strong selection pressures will exhibit high differentiation against a background of lower differentiation driven by unrestricted gene flow. Varying levels of differentiation across loci thus generates a mosaic of genomic differentiation. Methodological approaches designed to detect the processes underlying this mosaic, however, are confounded by demographic histories of secondary contact, genomic areas of suppressed recombination, recent divergences without gene flow, allele surfing, and selective sweeps specific to each lineage unrelated to the development of reproductive isolation (Noor & Bennett 2009; Cruickshank & Hahn 2014). For example, ecological speciation could result in genomic regions of elevated differentiation (i.e. genomic islands of differentiation) that are associated with niche partitioning, but not necessarily with reproductive isolation. Further, these islands of differentiation are mostly expected when adaptation occurs from moderate-to large-effect *de novo* mutations (Lackey & Boughman 2016). Genomic approaches used to identify islands of differentiation may thus be biased against identifying polygenic regions associated with species divergence, or towards identifying loci that contribute to ecological niche divergence of hybrids relative to both parental species and hence restrict gene flow between them. To avoid these biases, ecological niche divergence should be evaluated in the hybrid zone with respect to both parental species and correlated to patterns of hybrid ancestry (e.g. Hamilton *et al.* 2013).

Species of conifers are known to have ecologically differentiated niches despite the absence of strong morphological differences (e.g. Rehfeldt 1999). Strong pre- and post-zygotic isolating barriers contributing towards morphological disjunctions are often absent in conifers (Critchfield 1986; Buschiazzo *et al.* 2012; Pavy *et al.* 2012) due to their life history characteristics, such as longevity, high dispersal abilities, and long generation times (Petit & Hampe 2006; Neale & Kremer 2011). These contribute towards large effective population sizes, and moderate to high levels of genetic diversity, facilitating establishment across an array of ecological conditions. Ecological niche partitioning arising from extrinsic barriers is thus likely to play a dominant role in facilitating speciation within conifers (e.g. Hamilton *et al.* 2013).

In this study, we use an integrative approach to investigate processes leading to the divergence of two North American pine species – *Pinus strobiformis* Engelm. (southwestern white pine), and *P. flexilis* E. James. (limber pine). Our focal species inhabit a wide latitudinal range in the western part of North America, with a putative area of sympatry located in the southern Rocky Mountains and Colorado Plateau in which morphological evidence points towards the occurrence of hybridization (Steinhoff & Andresen 1971; Tomback & Achuff 2010). These species also display limited differences in morphological and reproductive traits (Benkman *et al.* 1984; Tomback *et al.* 2011; Bisbee 2014) and show evidence of local adaptation to the heterogeneous climatic conditions across their geographical range and also within the area of putative hybridization (Steinhoff & Andresen 1971; Moreno-Letelier *et al.* 2013; Borgman *et al.* 2015; Moreno-Letelier & Barraclough 2015; Goodrich *et al.* 2016). To examine the processes influencing species boundaries between these two conifer species, we asked three questions: (1) Is there niche divergence among *P. strobiformis*, *P. flexilis* and the putative hybrid zone? (2) Did the divergence of *P. strobiformis* and *P. flexilis* occur with continual gene flow? (3) Does a genome-wide mosaic of differentiation characterize divergence between *P. strobiformis* and *P. flexilis*, and is this pattern attributed to extrinsic, intrinsic, or an interaction of both factors? Our results are consistent with ecological divergence occurring with continual gene flow among the focal species, with several lines of evidence supporting the strong influence of extrinsic factors in reinforcing species boundaries.

## Materials and Methods

### Focal taxa and field sampling

*Pinus strobiformis* and *P. flexilis* are closely related species of white pines that occur across broad temperature and precipitation gradients in the mountainous areas of western North America. The native range of *P. strobiformis* includes Mexico and the southwestern United States, and its distribution exhibits disjunctions across dry and wet boreal mixed forest ecosystems (Looney & Waring 2013; Fig. 1). *Pinus flexilis* occurs in mountainous regions from northern Arizona and northern New Mexico to Alberta, with a region of putative sympatry with *P. strobiformis* in the southern Rocky Mountains and Colorado Plateau (Fig. 1). Across this zone of putative sympatry, cone morphology and dispersal syndromes fall along a continuum of divergence blending into the characteristics of populations in the allopatric zones of either species (Bisbee 2014).

**Fig. 1.**
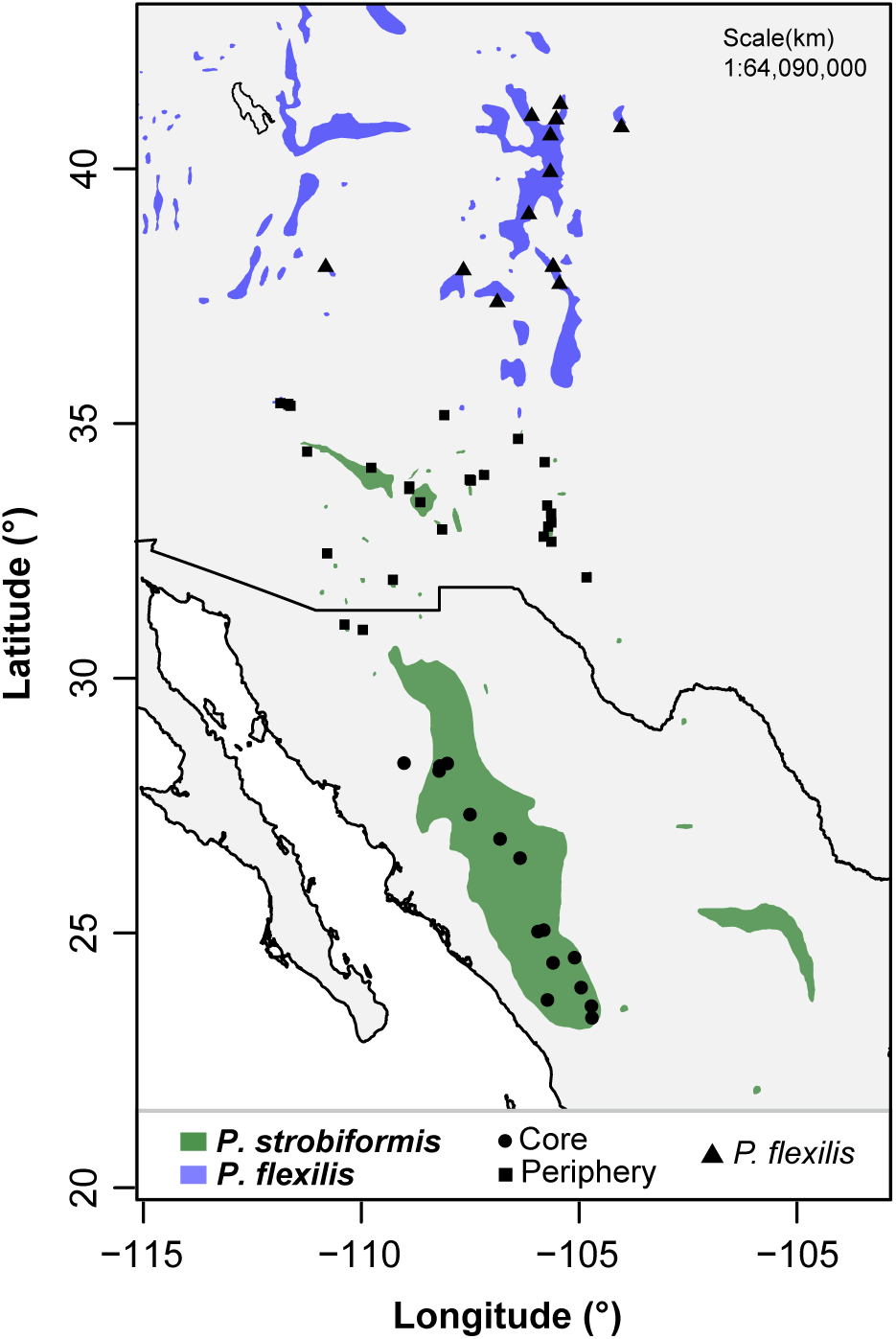
Map of sampling localities (black dots) overlaid on polygons showing geographical ranges of *Pinus strobiformis* (green) and *P. flexilis* (blue). The corresponding locality information is available in Table S1 of the Supporting Information.

We sampled 42 *P. strobiformis* populations encompassing a total of 376 trees (5-13 trees/population) from its entire geographical range. For *P. flexilis*, a total of 13 populations were sampled, with eight populations sampled from the southern periphery of the geographical range and five sampled closer to the range center (Fig. 1). Across these thirteen populations, we sampled a total of 69 trees (4–10 trees/population). To help minimize relatedness, trees within the same site were sampled with a minimum spacing of 50 m (*P. strobiformis*) and 200 m (*P. flexilis*) from each other.

### Data generation

#### Occurrence data

We assembled a comprehensive dataset of occurrences for ecological niche modeling (ENM) by supplementing our field site data with occurrence records downloaded from the Global Biodiversity Information Facility (GBIF) using functions from the DISMO package (Hijmans *et al.* 2017) available in the R environment (R Core Team 2017). Using a series of filtering steps to account for observation and sampling biases (Supporting Information, Appendix S1.A), we obtained a final dataset of 254 occurrence records for *P. strobiformis* and 420 for *P. flexilis*. Incorporating intraspecific genetic variation into ENMs can improve model fit and provide more robust predictions when projecting across time and space (Knowles *et al.* 2007; Ikeda *et al.* 2017). Thus, we divided presence locations within *P. strobiformis* into core (latitudinal range: 19–30.5 °N) and northern periphery (latitudinal range: 31–33°N). These groups likely represent different genetic clusters given the geographically restricted hybridization between *P. flexilis* and *P. strobiformis* (Steinhoff & Andresen 1971; Tomback & Achuff 2010; Bisbee 2014). We defined three groups that were the focus of our enquiries – (1) populations of *P. flexilis*, (2) populations of *P. strobiformis* from the northern range periphery (Periphery hereafter), and (3) populations of *P. strobiformis* from the range core (Core hereafter). Nineteen bioclimatic variables and altitude were used as predictors in the ENMs for all three groups. Present day geospatial data layers for these variables were downloaded from WorldClim v.1.4 (Hijmans *et al.* 2005) at 30 arc-second resolutions and at 2.5 arc-minute resolutions for the Last Glacial Maximum (LGM) and data were extracted from each layer using the RASTER package (Hijmans *et al.* 2016) available in R.

#### DNA sequence data

We extracted total genomic DNA from 445 individuals sampled across 55 populations of both species using DNeasy Plant Kits (Qiagen). Five ddRADseq libraries (Peterson *et al.* 2012), each containing up to 96 multiplexed samples, were prepared using the procedure detailed in Parchman *et al.* (2012). All libraries were digested using the *EcoR1* and *Mse1* restriction enzymes followed by ligation of adaptors, barcodes, and primers. Following PCR, we selected DNA fragments in the 300–400 bp size range using agarose gel electrophoresis followed by isolation of pooled DNA from these gels using QIAquick Gel Extraction Kits (Qiagen). Single-end sequencing, with one multiplexed library per lane, was used to obtain 100bp reads, with all sequencing conducted with Illumina HiSeq 2500 at the Nucleic Acids Research Facility located at Virginia Commonwealth University. The resulting FASTQ files were processed using dDocent bioinformatics pipeline (details in Supporting Information, Appendix S1.B; Puritz *et al.* 2014). The entire process yielded a total of 51 633 single nucleotide polymorphisms (SNPs), which were used as the starting dataset for all subsequent analyses.

### Data analysis

#### Ecological niche modeling and niche divergence

We developed ENMs for each of the three groups: Core, Periphery, and *P. flexilis*, using the algorithms available in Maximum Entropy (MaxEnt; Phillips *et al.* 2006). Since MaxEnt was specifically developed for presence-only data, we drew a one-degree rectangular buffer around the known distribution of both species and obtained 100 000 background points at random without duplicates in a cell. Data processing, model fitting, and model evaluation using 5,000 iterations within MaxEnt were conducted using the dismo, raster, rgdal (Bivand *et al.* 2017), and spThin (Aiello-Lammens *et al.* 2015) packages available in R. ENMs were constructed from climate variables with an absolute correlation coefficient (*r*) less than 0.85. Two indices were used to assess model performance for each group: overall regularized training gain (RTG) and area under the curve (AUC). Since LGM data were not available at 30 arc-seconds resolution, we built two ENMs for each group (2.5 arc-minutes and 30 arc-seconds), but only used the 2.5 arc-minutes models for hindcasting to infer historical patterns of sympatry between species that could facilitate gene flow. We followed an average projection ensemble approach across three LGM scenarios (CCSM4, MIROC & MPI) to obtain a hindcasted suitability map. Changes in habitat suitability (stability) were assessed by adding MaxEnt-predicted suitability maps across the LGM and present (as in Ortego *et al.* 2015). For these maps, values closer to 2 in a gridded cell are associated with the stability of highly suitable habitat for a given group across time points. In contrast, values closer to 0 are associated with the stability of highly unsuitable habitat for a given group across time points. Suitability scores obtained across the full geographical extent for the present conditions at 30 arc-seconds were compared across all three pairs of groups (Core–Periphery, Core–*P. flexilis*, Periphery–*P. flexilis*) to investigate patterns of niche evolution. To account for potential biases towards niche divergence introduced by latitudinally associated environmental variation in the present range of each pair, we performed an asymmetric background randomization test, based on Schoener’s *D*, in the R package ENMTools (Warren *et al.* 2008). The two resulting null distributions of niche divergence obtained through this test correspond to the background of each group compared against the other. An observed value of Schoener’s *D* much smaller than expected after accounting for background differences is indicative of niche divergence, whereas a value much larger than expected indicates niche conservatism (Warren *et al.* 2008).

#### Population structure and demographic modeling

We assessed the pattern and the extent of genetic divergence between *P. strobiformis* and *P. flexilis* using multiple methods. First, we grouped the 42 *P. strobiformis* populations into the same core and periphery groups discussed above (see Data Generation & Fig. 1). We conducted principal components analysis (PCA) to visualize grouping of sampled trees into the three groups delineated in our methods (Patterson *et al.* 2006; McVean 2009). To complement the PCA, we also conducted an individual-based assignment test using fastSTRUCTURE, a variational Bayesian version of STRUCTURE designed for use with large SNP datasets (Raj *et al.* 2014). For any given number of predefined clusters (*K*), FastSTRUCTURE assigns a *Q*-value representing the proportion of a sample’s ancestry derived from each cluster. We set *K* to 2, representing the two parental species investigated here, as we were interested in admixture between two defined species and not the potential number of groups within our genetic data. Lastly, we utilized hierarchical fixation indices (*F*-statistics) to assess the extent of differentiation between species by nesting trees into populations and populations into species. There are two levels within the hierarchy, with *F*_*CT*_ describing differentiation among groups at the highest level of the hierarchy and *F*_*ST*_ describing differentiation among groups across all levels of the hierarchy (see Yang 1998). A similar nested model with the highest level of hierarchy being groups within *P. strobiformis* was used to assess intraspecific differentiation. For the former, *F*-statistics are denoted using the term ‘species’ in the subscripts, whereas the latter uses the term ‘groups’ in the subscripts. We used a similar hierarchical model with variance partitioning to estimate group specific and pairwise *F*-statistics for the three groups delineated in this study. We denote pairwise values of *F*_*ST*_ using one-letter abbreviations for the groups being compared (e.g. *F*_*ST*-CP_ indicates *F*_*ST*_ between Core and Periphery), and group specific values of *F*_*ST*_ with the name of the group in subscripts. We constructed 95% confidence intervals of multilocus *F*-statistics using bootstrap resampling (*n* = 100 replicates) in the HIERFSTAT package (Goudet 2005) available in R. Along with estimation of *F*-statistics, we also assessed overall levels of genetic diversity using multilocus estimates (i.e. means across SNPs) of observed and expected heterozygosities (*H*_o_ and *H*_e_) per population.

Presence of individuals with mixed ancestry, as identified using fastSTRUCTURE, can be a result of secondary contact, recent divergence causing incomplete lineage sorting, or the presence of gene flow throughout the divergence history. Disentangling these explanations is important, because it directly influences our understanding of the relative importance of intrinsic and extrinsic factors in facilitating speciation. For instance, when speciation is recent or has occurred with gene flow, we expect a heterogeneous landscape of genomic differentiation, such that while most of the genome is freely introgressed between species, only a few genomic regions associated with intrinsic or extrinsic factors inhibiting gene flow will exhibit elevated differentiation (Wu 2001; Feder *et al.* 2012). However, if hybrids are formed in areas with novel habitats, introgression might be selectively advantageous, such that heterozygotes at climate-associated loci will confer higher fitness, and will lack elevated islands of differentiation. To infer the timing and influence of various demographic processes shaping the divergence history of our focal groups, we conducted demographic modeling using Diffusion Approximation for Demographic Inference (∂A∂I; Gutenkunst *et al.* 2009).We down-sampled the total SNP dataset for computational simplicity based on population genetic summary statistics and then randomly sampled one SNP per assembled contig to obtain a final dataset of 4,800 SNPs that were used in subsequent ∂A∂I analyses (Supporting Information, Appendix S1.C).

We compared a model of pure divergence with no gene flow (*M*_*1*_) against a set of 10 alternative demographic models (*M*_*2*_–*M*_*7*_) representing different speciation scenarios including varying timing and directionality of ancient or contemporary gene flow (Supporting Information, Fig. S1). Complexity was added to the models with gene flow by incorporating heterogeneity in the gene flow parameter across loci (Tine *et al.* 2014, models *M*_*8*_–*M*_*11*_, Fig. S1). We ran 10 replicate runs of each model in ∂A∂I, using a 200 × 220 × 240 grid space and the nonlinear Broyden-Fletcher-Goldfarb-Shannon (BFGS) optimization routine. Following Carstens *et al.* (2013), we conducted model selection in an information-theoretic framework using Akaike information criterion (AIC; Akaike 1974) and ΔAIC (AIC_model i_ - AIC_best model_) scores (Burnham & Anderson 2002), calculated using results from the best replicate run (highest composite likelihood) for each model. Unscaled parameter estimates were obtained using a per-site substitution rate of 7.28 × 10^−10^ substitutions/site/year rate estimated for Pinaceae by De La Torre *et al.* (2017) and a generation time of 50 years.

#### Genomics of interspecific introgression

Analyses of clines across hybrid zones are widely used to identify loci exhibiting exceptional patterns of introgression relative to the average genomic background (Fitzpatrick 2013; Gompert *et al.* 2012a; Gompert & Buerkle 2011; Stankowski *et al.* 2015). We classified our sampled trees into categories corresponding to admixed (*n*_A_ = 111) and parental species (*P. strobiformis* = 277, *P. flexilis* = 54) based on the *Q*-values from fastSTRUCTURE. Trees with *Q*-values of 0.9 or higher were classified as pure *P. strobiformis*, those with *Q* of 0.1 or lower were classified as pure *P. flexilis*, and those with intermediate *Q*-values were classified as admixed (e.g. Ortego *et al.* 2014). As most loci exhibited little to no differentiation between parental species, we retained only loci with a minor allele frequency (MAF) difference of at least 10% between parental species (*n* = 4,857 SNPs). This allowed us to avoid false correlations between cline parameters and fixation indices (Parchman *et al.* 2013). We used this subset of 4,857 SNPs to perform a Bayesian genomic cline analysis in BGC v1.0 (Gompert & Buerkle 2012; Gompert & Buerkle 2011). Using Markov chain Monte Carlo (MCMC) sampling, BGC estimates the posterior distribution of ancestry for each locus as a function of the genome-wide admixture coefficient. The BGC model includes two genomic cline parameters, *α* (genomic cline center) and *β* (genomic cline rate, i.e. slope), determining the probability of *P. flexilis* ancestry, and the rate of transition from *P. flexilis* to *P. strobiformis* given a level of genomic admixture described by the hybrid index, *h*, respectively (Gompert & Buerkle 2012; Gompert *et al.* 2012a). A tree with *h* = 0 was classified as having solely *P. strobiformis* ancestry, whereas a tree with *h* = 1 was classified as having solely *P. flexilis* ancestry. We ran BGC for five replicate runs, each 45 000 steps in length, and, after discarding the first 25 000 steps as burn-in, we thinned the posterior distribution every 20 steps, thus yielding 1,000 samples which were used for inference of model parameters. We used TRACER v1.6 (Rambaut *et al.* 2013) to test for convergence among replicated runs, as well as appropriate mixing along MCMC chains. We identified loci with excess ancestry (relative to the genome-wide average) as those with posterior *α* or *β* credible intervals (CI; 95% equal-tail intervals) not containing zero. Moreover, we identified outlier loci as those with posterior mean point estimates of *α*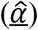 or *β*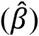 significantly different from the rest of the genome, as judged by comparison to posterior quantiles of random-effect priors for *α* and *β* (Gompert *et al.* 2012a). Several empirical and simulation based studies have demonstrated that both *α* and *β* can reflect patterns of selection in the hybrid zone (Gompert et al. 2012b; Janoušek *et al.* 2012), but the interpretation of these values is influenced by the underlying demographic scenario (Gompert & Buerkle 2012; Gompert *et al.* 2012a,2012b). Besides categorizing loci, we also tested for correlations among locus-specific *F*_*CT-species*_, *α*, and *β*, with and without absolute values for *α* and *β*. The sign of the cline parameters (specifically *β*) have direct implications for inferring the processes maintaining species boundaries and hence were incorporated in correlation tests. Specifically, extremely positive values of *β* reflect strong selection against hybrids or population structure in the hybrid zone (Gompert *et al.* 2012b), while extremely negative values of *β* indicate a wide cline representing easy dispersal across species boundaries (Janoušek *et al.* 2012).

Although the hybrid index (*h*) obtained from BGC provides information about the age and stability of a hybrid zone, such inferences are limited to only one generation of admixture (Fitzpatrick 2012). We estimated *h* and interspecific heterozygosity using INTROGRESS (Gompert & Buerkle 2010), in order to extend our interpretations to a historical hybrid zone and categorize individuals into recent (F1s), advanced generation (FNs), and backcrossed hybrids (BC). This was done using a modified classification from Hamilton *et al.* (2013). Both BGC and INTROGRESS yielded very similar estimates of *h* (Pearson’s *r* = 0.7, p = 0.00042), thus we used estimates from INTROGRESS due to the availability of inter-specific heterozygosity estimates from this software. To test for the influence of exogenous factors in the maintenance of species boundaries we performed linear regression analyses with backward variable selection using *h* against climate and geography as predictor variables.

## Results

### Ecological niche modeling and niche divergence

ENMs for each of the three groups used in this study (Fig. 1) had high predictive ability, as indicated by AUC and RTG values (Table 1). For Core and Periphery, several covariates stood out as important with precipitation seasonality (Bio15) being shared between Core and Periphery; however, altitude was consistently the most important variable for *P. flexilis* across different measures of variable importance (Table 1). Hindcasting the 2.5 arc-minute model onto LGM data layers supported a recent, post-LGM niche fragmentation and northward expansion in Periphery (Supporting Information Fig. S2). A similar post-LGM northward expansion of suitable niche space was observed for *P. flexilis*. Furthermore, there was extensive range overlap between the two species during the LGM, which was greater than what is currently observed (Supporting Information Fig. S2). The values of niche similarity based on Schoener’s *D* ranged from 0.05 (*P. flexilis*–Core) to 0.17 (Periphery – Core). Background randomization tests revealed statistically significant niche divergence for two of the three comparisons (Fig. 2). Specifically, there was asymmetrical niche divergence between Core and Periphery, with the niche of periphery being conserved relative to the background of core. A similar pattern was noted using only the presence points, where each group formed a distinct cluster within the multivariate climate space defined by the top two principal components (PCs) derived from PCA on the climate variables used for construction of the ENMs (Supporting information, Fig. S3.A).

**Table 1.** Ecological niche model performance and variable importance at 30 arc-second resolution

| Groups | AUC | RTG | RTG importance <sup>#</sup> | Permutation importance <sup>¢</sup> | Percent contribution <sup>¢</sup> | Regression coefficient importance <sup>*</sup> |
| --- | --- | --- | --- | --- | --- | --- |
| Core | 0.97 | 2.51 | Bio15<sup>1</sup>, Bio4<sup>2</sup> | Bio4 | Altitude | Bio4 |
| Periphery | 0.99 | 3.92 | Bio9<sup>3</sup> | Bio10<sup>4</sup>, Bio9, Bio6<sup>5</sup> | Altitude | Bio15 |
| *P. flexilis* | 0.94 | 1.72 | Altitude | Altitude | Altitude | Altitude |

AUC: Area under the curve; RTG: Regularized training gain

**Fig. 2.**
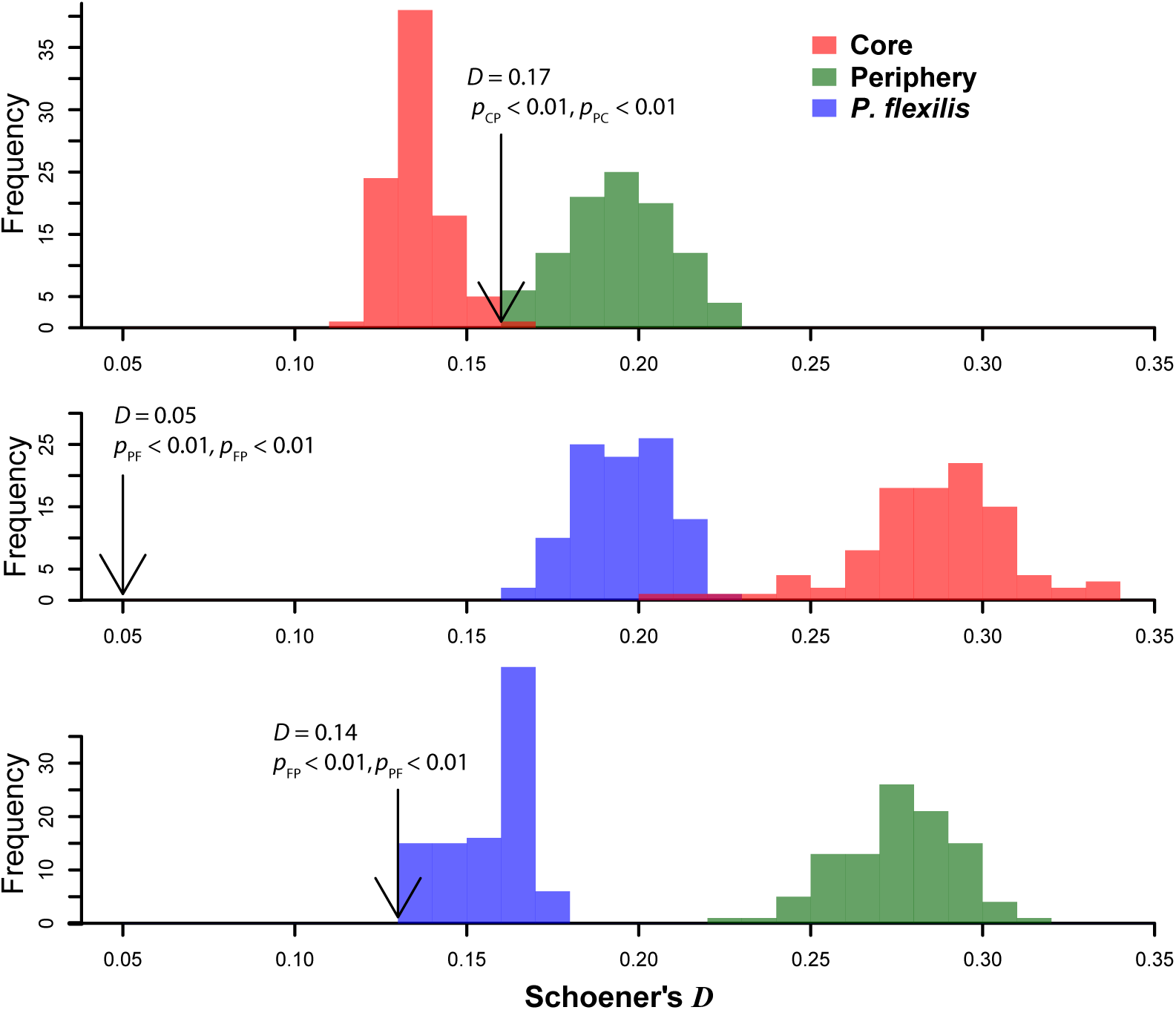
Results of niche divergence tests (Schoener’s *D*) for all pairwise comparisons among groups. Histograms indicate the background levels of niche divergence and arrows indicate the observed value of Schoener’s *D* for each pair compared.

### Population structure and divergence history

The PCA using 51 633 SNPs was consistent with trees sampled from Core being differentiated from those of *P. flexilis*, which was most marked along PC1 (Fig. 3A). This PC explained 0.9% of the total genetic variance, which was in line with the overall level of differentiation estimated using hierarchical *F*-statistics (*F*_*ST*-species_ = 0.021, 95% CI: 0.008–0.031). Trees sampled from Periphery were located between those sampled from Core and *P. flexilis* (Fig. 3A), in line with peripheral populations containing hybrids between the two parental species. There was also a latitudinal gradient in the mean population *Q*-values, as estimated using fastSTRUCTURE, with Core populations exhibiting little to no ancestry from *P. flexilis* and Periphery being a mixture of *P. flexilis* and Core (Fig. 3B). At the individual tree level, we observed a strong negative correlation (Pearson’s *r* = −0.69, *p* = 4.087e-07) between *Q-*values of putative hybrids and latitude, which is consistent with a geographical gradient of genomic introgression where trees geographically proximal to either parental species contain more ancestry from that parental species. Multilocus estimation of differentiation between species (*F*_CT-species_) was 0.01 (95% CI: 0.005–0.018, Fig. 5A), while that between groups within *P. strobiformis* (*F*_CT-groups_) was 0.003 (95% CI: 0.0007–0.006). Multilocus *F*_ST_ within each group, pairwise *F*_*ST*_ between each pair of group, and heterozygosities differed little among the three groups, with the Core–*P. flexilis* comparison having the highest pairwise *F*_*ST*-CF_ = 0.019 (Table 2). Although populations of Periphery exhibited slightly higher heterozygosities and *F*_*ST*_ values (*F*_*ST*-periphery_), this pattern was mainly driven by few populations, as indicated by the wider confidence interval around these estimates (Table 2).

**Fig. 3.**
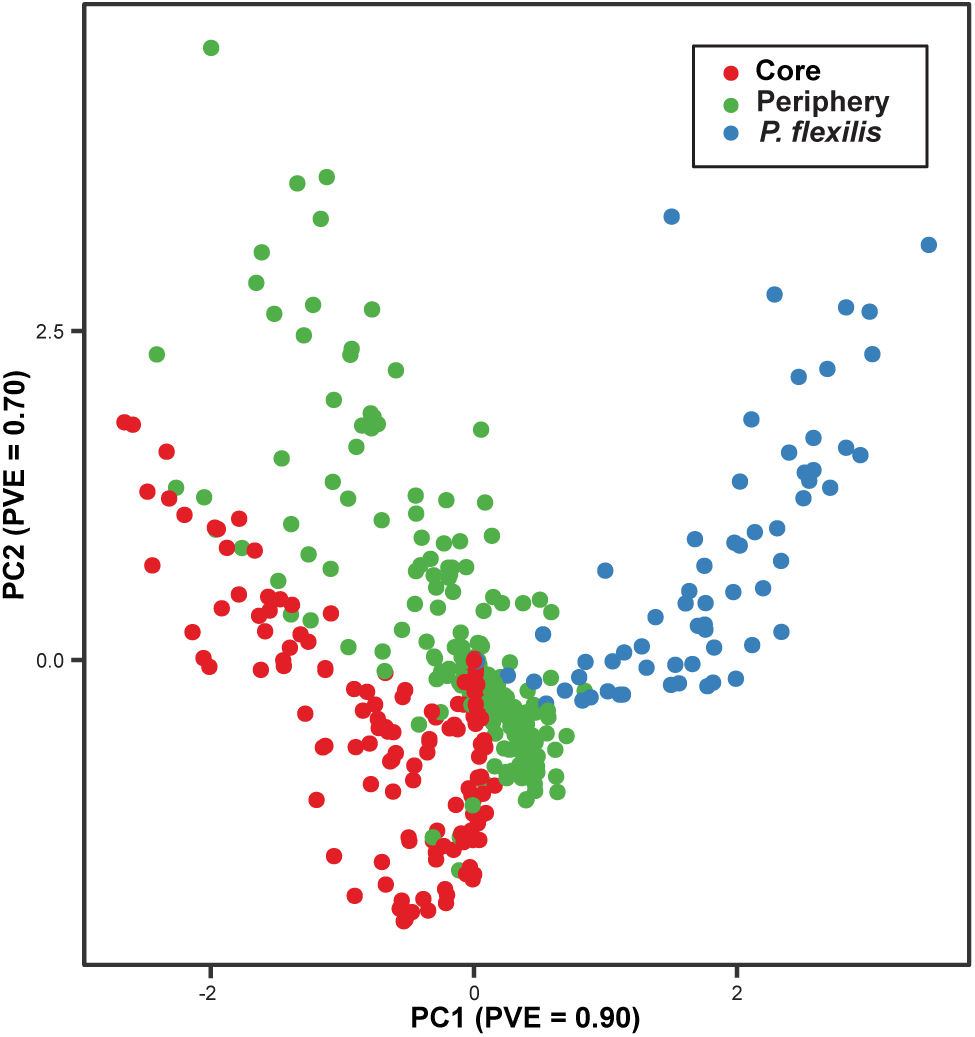
**A**) Results of population genetic structure analysis using PCA on 51 633 SNPs. **B)** Results of assignment analyses for each tree in fastSTRUCTURE for *K* = 2 clusters (right panel) plotted onto a topographic map of the study area (left panel). Each pie chart represents the average ancestry of a population from *P. strobiformis* and *P. flexilis*.

**Table 2.** Estimates of genetic diversity and divergence within and across the three groups, compared to a genome-wide *F*_*ST-species*_ of 0.02 (95% CI: 0.008–0.03) and *F*_*ST*-strobiformis_ of 0.009 (95% CI: 0.007–0.014).

| Group | Multilocus $F_{ST}$<br>(95% CI) | Pairwise $F_{ST}$<br>(95% CI) | Mean $H_e \pm$ s.d. | Mean $H_o \pm$ s.d. |
| --- | --- | --- | --- | --- |
| Core | 0.003 (0.0025–0.0034) | periphery: 0.009(0.001–0.023)<br>*P. flexilis*: 0.019 (0.006–0.032) | 0.135 ± 0.01 | 0.111 ± 0.01 |
| Periphery | 0.007(0.0071–0.0073) | *P. flexilis*: 0.015 (0.005–0.024)<br>core: 0.009(0.001–0.023) | 0.133 ± 0.02 | 0.105 ± 0.03 |
| *P. flexilis* | 0.003 (0.0025 – 0.0041) | core: 0.019 (0.006–0.032)<br>periphery: 0.015 (0.005–0.024) | 0.130 ± 0.01 | 0.111 ± 0.01 |

The best-supported demographic model was *M*_*4*_, which is a model of symmetric ancient gene flow between ancestral *P. strobiformis* and *P. flexilis* lineages, followed by contemporary gene flow between Periphery and *P. flexilis* (Table 3; Fig 4). This model was supported by a distinct minimum AIC score that was better than that of all other ∂A∂I models by a margin of 44.8 information units (ΔAIC_*i*_ = 44.8 or greater). The next best model, *M*_*8*_, was similar to that of *M*_*4*_, but without contemporary gene flow between *P. strobiformis* and *P. flexilis*, and a heterogeneous ancient gene flow between ancestral populations of the two parental species. Converted parameter estimates indicated that the species diverged 11.36 million years ago (Ma) in the Miocene, but that groups within *P. strobiformis* diverged 2.29 Ma during early Pleistocene (Fig. 4). Overall rates of gene flow between species were substantial for both historical and contemporary periods, however the contemporary gene flow between species was geographically restricted Periphery (Supporting Information Table S2). In addition, *P. flexilis* and Periphery experienced asymmetrical gene flow, which was larger in the direction of *P. flexilis* to Periphery (*M*_*PF*_ = 8.81 migrants/generation versus *M*_*FP*_ = 4.35). Periphery had the largest population size estimate, while *P. flexilis* was inferred to have experienced a reduction in population size through time.

**Fig. 4.**
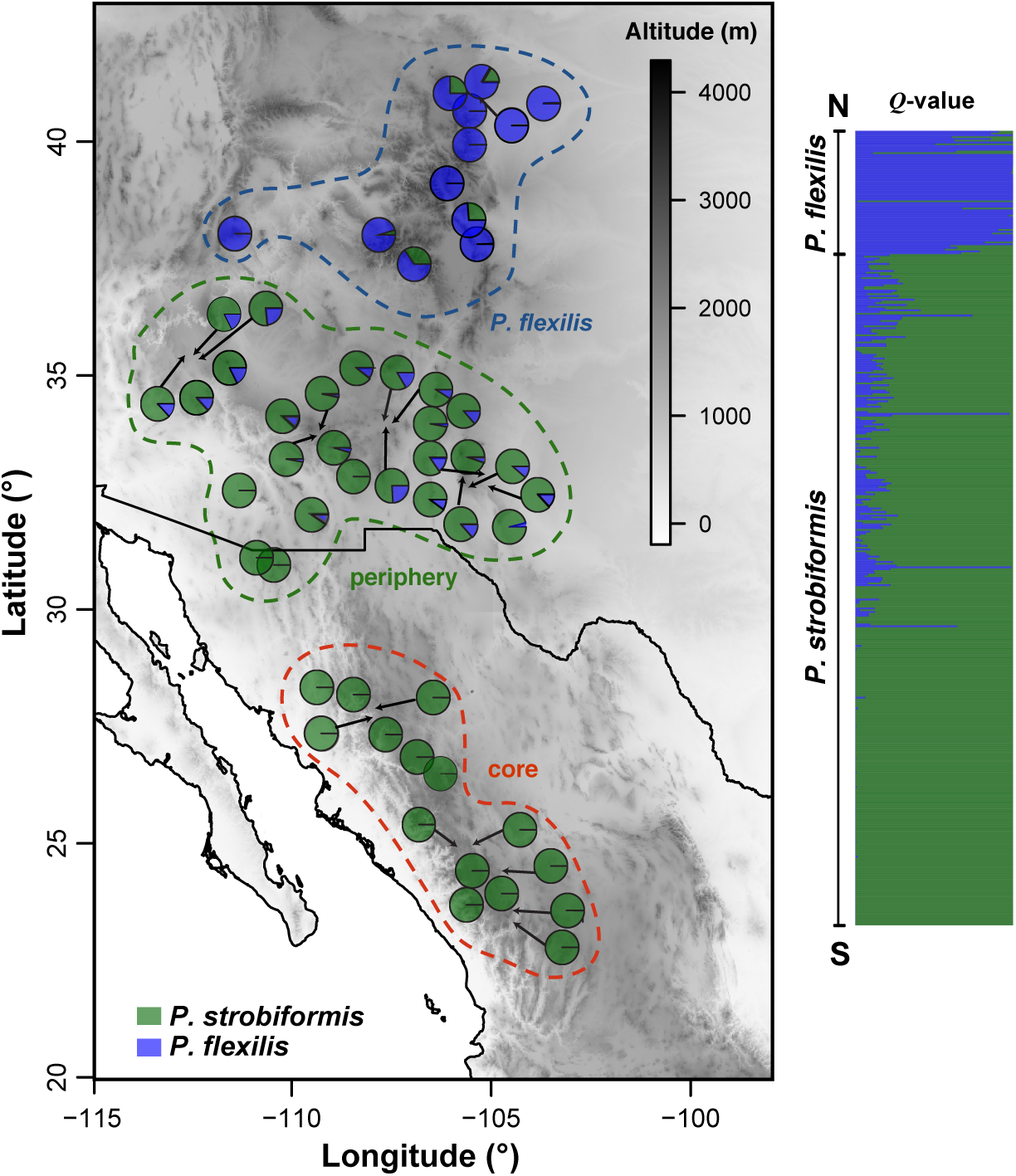
The best-supported model from ∂A∂I analysis. This figure shows the optimized parameter estimates for divergence times (*T*_i_) in units of millions of years ago (Ma), reference effective population size (theta; or after conversion, *N*_eref_), lineage population sizes (*N*_i_), and rates of gene flow (*M*_ij_) for the optimal model determined by AIC model selection (see results in Table 2).

**Table 3.** Model composite likelihoods and AIC model selection results for 11 alternative demographic models of *P. strobiformis* (core and periphery)–*P. flexilis* divergence. The best supported model, that with the minimum AIC score (hence, ΔAIC_*i*_= 0), is underlined, and the two best models are shown in boldface.

| Model | Model description | $\ln$ Composite likelihood | $k$ | AIC | $\Delta\text{AIC}_i$ |
| --- | --- | --- | --- | --- | --- |
| $M_1$ | Strict isolation, no gene flow | –883.143112 | 6 | 1778.29 | 65.44 |
| $M_2$ | Secondary contact (Periphery–*P. flexilis*) | –886.227416 | 7 | 1786.45 | 73.60 |
| $M_3$ | Ancient gene flow (speciation with gene flow) | –888.003307 | 7 | 1790.01 | 77.16 |
| **<u>$M_4$</u>** | **<u>Ancient gene flow, plus Periphery–*P. flexilis* gene flow</u>** | **<u>–847.424540</u>** | **<u>9</u>** | **<u>1712.85</u>** | **<u>0.00</u>** |
| $M_5$ | Ancient gene flow, plus Core–periphery gene flow | –885.428135 | 9 | 1788.86 | 76.01 |
| $M_6$ | Secondary contact (Periphery–*P. flexilis*) and Core–Periphery gene flow | –883.949484 | 10 | 1787.90 | 75.05 |
| $M_7$ | Ancient gene flow, followed by Periphery–*P. flexilis* gene flow, and Core–Periphery gene flow | –892.210862 | 9 | 1806.42 | 93.57 |
| **$M_8$** | **Heterogeneous ancient gene flow** | **–869.824520** | **14** | **1757.65** | **44.80** |
| $M_9$ | Heterogeneous ancient gene flow, plus Core–Periphery gene flow | –884.511096 | 11 | 1791.02 | 78.17 |
| $M_{10}$ | Heterogeneous gene flow during secondary contact (Periphery–*P. flexilis*), and Core–Periphery gene flow | –902.279445 | 9 | 1828.56 | 115.71 |
| $M_{11}$ | Heterogeneous ancient gene flow, followed by heterogeneous gene flow between Periphery–*P. flexilis*, and between Core–Periphery | –922.814525 | 11 | 1873.63 | 160.78 |

AIC, Akaike information criterion; $k$ , the number of parameters in the model; $\ln$ , natural logarithm.

### Genomics of interspecific introgression

Values of *h* ranged from near zero to 0.80, with values around 0.20 being the most common thus suggesting overrepresentation of *P. strobiformis* ancestry (Fig. 5B). Estimates of interspecific heterozygosity had a narrow range from 0.45 to 0.64, indicating weak reproductive barriers (Hamilton *et al.* 2013) and a long history of recombination within the hybrid zone (Gompert *et al.* 2014). Classification of trees into genotypic classes based on *h* and interspecific heterozygosity revealed a dominance of advanced-generation hybrids (54%), with some trees being backcrossed into *P. strobiformis* (22%), but no recent hybrids (F1s) were apparent. Stepwise linear regression analysis revealed a significant effect of geography and climate on *h* across the putative hybrid zone. Latitude (Pearson’s *r* = 0.41, *p* = 4.087e-07), precipitation seasonality (Pearson’s *r* = −0.32, *p* = 0.006), and mean temperature of the warmest quarter (Pearson’s *r* = −0.18, *p* = 0.005) had a strong influence on *h*, in line with the latter two being important predictor variables for Periphery in our ENM.

**Fig. 5.**
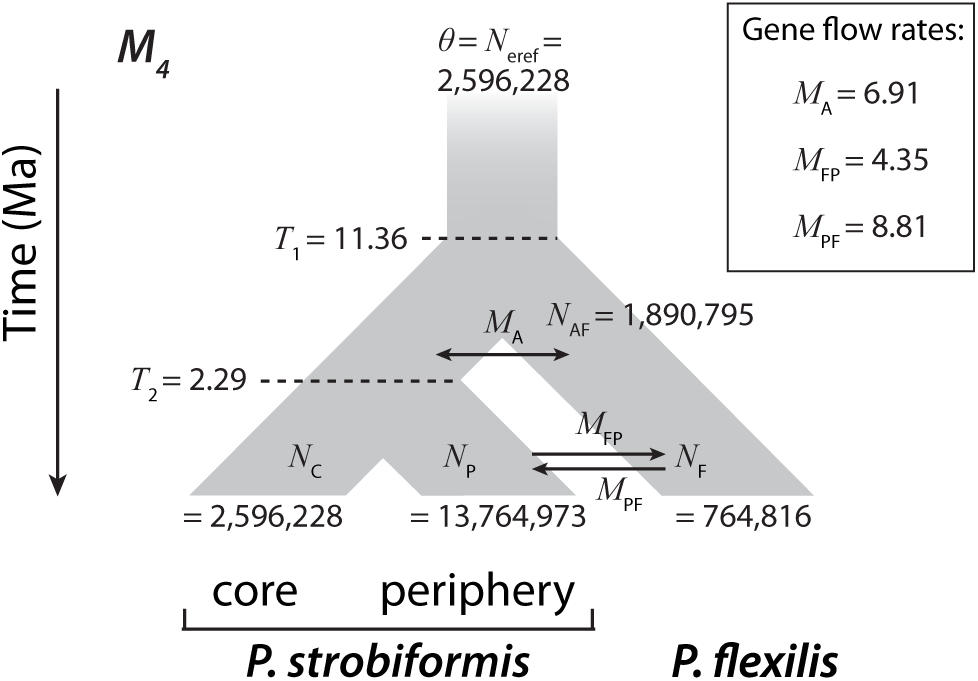
**A**) Genomic distribution of *F*_CT_ **B)** frequency distribution of hybrid index **C)** variation in genomic ancestry as a function of hybrid index **D)** correlation between genomic cline parameters, and **E)** 3D correlation plot of genomic cline parameters and *F*_*CT*_

The influence of selection is reflected in both *α* and *β*; however, exceptional values of *α* are more indicative of selection favoring directional introgression, while extreme *β* values reflect strong selection against hybrids or presence of population structure within the hybrid zone (Gompert *et al.* 2011). Substantial variation was found in estimates of genomic cline parameters (Fig. 5D,C), especially for *α*, with its range (-0.99 to 1.72) being 18.5-fold as wide as that of the range for *β* (-0.068 to 0.078). Similar to the patterns observed in the distribution of *h*, an asymmetry towards *P. strobiformis* ancestry was noted in the genomic cline estimates. From the posterior distribution of *α*, we found 3,193 outlier loci, of which 570 (17.9%) had elevated probabilities of *P. flexilis* ancestry (positive 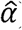), and 2,623 (82.1%) had elevated probabilities of *P. strobiformis* ancestry (negative 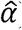). We identified fewer loci with excess ancestry, but in contrast to the pattern for outlier loci those with excess ancestry favored *P. flexilis* over *P. strobiformis* ancestry. Among the 287 loci with excess ancestry, 204 (71.1%) had excess *P. flexilis* ancestry (i.e. lower 95% CI of *α* > 0) and 83 (28.9%) had excess *P. strobiformis* ancestry (i.e. upper 95% CI of *α* < 0). The multilocus *F*_*CT*-species_ estimate for loci with excess ancestry was 0.12 (95% CI: 0.09-0.13) while for outlier loci it was 0.058 (95% CI: 0.05-0.09). We did not identify any loci that were *β* outliers or had excess ancestry indicated by *β*. Hierarchical *F*_CT-species_ between species was negatively correlated with raw values of *α*(Pearson’s r =-0.036, *p* = 0.01), positively with raw values of *β* (Pearson’s r = 0.048, *p* = 0.0007) and positively with absolute values of both *α*(Pearson’s r = 0.14, *p* = 2.2e-16) and *β*((Pearson’s r = 0.26, *p* = 2.2e-16) (Fig. 5E).

## Discussion

We identified strong evidence supporting ecological divergence between *P. strobiformis* and *P. flexilis* despite extensive gene flow. Our findings are generally consistent with previous reports on the species examined here; however, in contrast to the recent divergence time estimates arrived at using chloroplast and mitochondrial loci (Moreno-Letelier *et al.* 2013), our demographic modeling results reject a hypothesis of extremely recent divergence between the two species. Instead, we support a model of ongoing speciation with gene flow that is driven and maintained primarily by extrinsic factors.

### Niche evolution and ecological divergence

Our results indicate that climatic factors have played a major role in driving niche divergence between *P. strobiformis* and *P. flexilis*. Populations within Periphery coincide with the known hybrid zone between *P. strobiformis* and *P. flexilis* (Steinhoff & Andresen 1971; Tomback & Achuff 2010; Bisbee 2014) and formed a distinct group characterized by niche divergence from *P. flexilis* and asymmetrical niche divergence from Core. The asymmetrical pattern of niche divergence between Core and Periphery is likely a result of recent divergence. Under this scenario, we expect that niche differentiation would occur primarily along a few environmental variables that strongly influence fitness in the transitional environmental conditions, with little to no differentiation among groups on the other environmental axes. For example, precipitation seasonality was an important niche predictor for both Core and Periphery, although they were differentiated along this environmental axis (Supporting information, Fig. S3.B). This pattern reiterates the presence of hybrid populations in transitional environmental conditions and experiencing early stages of niche divergence from both parents.

In line with these results, precipitation seasonality and mean temperature of warmest quarter had a strong negative association with genomic ancestry and contributed to the niche divergence of Periphery. These two climatic variables influence plant evapotranspiration and thus affect drought responses (Mishra & Singh 2010). Drought stress during the active growing season is widely recognized as a limiting factor to plant growth in the western parts of North America (Williams *et al.* 2010; Restaino *et al.* 2016) and our results are indicative of adaptive divergence along a drought tolerance gradient between the three groups (Gitlin *et al* 2006; Allen & Breshears 1998). Further, our study broadly agrees with other reports in *P. strobiformis* indicating precipitation and altitude to be some key niche predictors (Aguirre-Gutiérrez *et al.* 2015; Shirk *et al.* 2017). Climatic clines of admixture and environmentally-dependent maintenance of hybrid zones have been noted in other species of woody perennials in the genera *Quercus* (Dodd & Afzal-Rafii 2004), *Picea* (Hamilton *et al.* 2013; De La Torre *et al.* 2014b), *Rhododendron* (Milne *et al.* 2003), and *Pinus* (Cullingham *et al.* 2014). An important variable unaccounted for in our study, however, is white pine blister rust infestation (*Cronartium ribicola* Fisch J.C), which influences stand dynamics of both species (Looney & Waring 2013) and drives trade-offs between drought-tolerance and resistance in *P. flexilis* (Vogan & Schoettle 2015). However, given the low and recent incidence of infestation (Looney *et al.* 2015), it is unlikely that *C. ribicola* contributed towards niche divergence of the focal taxa. It is more likely that under projected scenarios of environmental change, interactions between these selective forces may occur in the future to influence ongoing speciation dynamics.

Despite fluctuations in suitable range size (Supporting information, Fig. S2) and previous studies indicating reduction in genetic diversity at range margins using chloroplast markers (Moreno-Letelier & Piñero 2009), we find no evidence for this in our study. This might be explained by the asymmetry in gene flow between Periphery and *P. flexilis*, as inferred from our demographic modeling results (Bridle & Vines 2007; Ortego *et al.* 2014). Moreover, evidence of directional introgression from *P. flexilis* (positive *α* outliers) might also have facilitated adaptation to transitional environmental conditions. Such novel allelic combinations have often contributed to the ability of populations to colonize new niches that are intermediate to the climatic conditions experienced by the parental species (De Carvalho *et al.* 2010; Hamilton *et al.* 2013; De La Torre *et al.* 2014b; Geraldes *et al.* 2014). Presence of a locally adapted and historical hybrid zone is supported by the absence of *β* outliers in our genomic cline results (Kamdem *et al.* 2016) and a recent study uncovering high *Q*_*ST*_ values associated with physiological traits primarily linked to drought tolerance within the group Periphery (Goodrich *et al.* 2016). The geographic cline in *h*, asymmetry in excess ancestry loci towards *P. flexilis*, and elevated estimates of *F*_*ST*–periphery_, however, indicate the potential for geographically driven neutral introgression to generate biased signals of local adaptation within the peripheral populations (Geraldes *et al.* 2014). Ongoing investigations using replicate populations in the hybrid zone across gradients of geographic proximity and climate similarity will be able to address this issue in further detail (Lotterhos & Whitlock 2015; Riquet *et al.* 2017).

### Speciation with gene flow without islands of divergence

Demographic modeling indicated that divergence of *P. strobiformis* and *P. flexilis* is not recent (~11 Ma) on an absolute time scale and has occurred with continuous gene flow. The presence of continual gene flow and absence of a period of allopatry, moreover, is also supported by the L-shaped distribution of *F*_CT-species_ values (Fig. 5A. Nosil & Feder 2012). Reduction in overlapping niche suitability from LGM to present, between *P. strobiformis* and *P. flexilis*, agrees with the best-supported demographic model indicating continuous and geographically restricted contemporary gene flow (also see Moreno-Letelier & Piñero 2009). Contemporary reduction in *N*_*e*_ for *P. flexilis* from our demographic modeling is contrary to the predicted post-LGM expansion of suitable habitat. This is likely due to the limited geographical sampling within *P. flexilis* for our genomic analyses, the two modeling approaches estimating population sizes across very different temporal scales, and a nonlinear relationship between habitat suitability and realized population sizes. Despite the potential for islands of divergence under a model of speciation with gene flow (Nosil 2008; Feder *et al.* 2012; Tine *et al.* 2014), as well as niche divergence results consistent with ecological speciation with gene flow between *P. strobiformis* and *P. flexilis*, the best-supported demographic model herein did not provide evidence for islands of divergence.

The absence of elevated islands of divergence in this study, however, does not necessarily indicate an absence of adaptive divergence during speciation with gene flow. The lack of islands of divergence is expected in conifers, given the prevalence of polygenic architectures defining continuous trait variation across species boundaries and the expected prevalence of soft sweeps (Pritchard & Rienzo 2010; Alberto *et al.* 2013; Rajora *et al.* 2016; Lind *et al.* 2017). Alternatively, given the large and complex genomes of conifers (reviewed by De La Torre *et al.* 2014a) our ddRADseq markers underrepresented genic regions, which are often identified as islands of divergence (Nosil & Feder 2012; Zhou *et al.* 2014; Moreno-Letelier & Barraclough 2015; Marques *et al.* 2017). For example, Moreno-Letelier & Barraclough (2015) demonstrated the potential for islands of divergence at drought-associated genes, which had a high average *F*_*ST*_ value of 0.33 (0.09–0.4) compared to the genome-wide estimate from this study (*F*_ST-species_ = 0.02). Future investigations using candidate gene approaches or exome capture might thus be able to identify islands of divergence in conifers, although evidence of adaptation in complex genomes often also appears within intergenic regions (Li *et al.* 2012).

### Genomic mosaic of introgression

The spatial context of loci within genomes, as well as the temporal scale of divergence between lineages, can influence patterns of introgression and are often depicted by a mosaic landscape of genomic differentiation and ancestry. For instance, Coyne & Orr (1989), Noor & Bennett (2009), and Christe *et al.* (2017) have all argued that islands of divergence tend to accumulate around regions of reduced recombination such as centromeres and inversions. Extrinsic factors, such as disruptive selection can also restrict gene flow, but under the observed demographic scenario these alone are unlikely to generate islands of divergence (Yeaman & Otto 2011; Yeaman *et al.* 2016). However, extrinsic barriers can often result in the evolution of intrinsic barriers and subsequently become coupled with intrinsic barriers and with several other loci experiencing similar selection pressures (Agrawal *et al.* 2011; Flaxman *et al.* 2014). Thus, given sufficient time, even under a model of speciation with gene flow, such coupling effects will ensure the maintenance of species boundaries relative to the action of either factor alone (Barton & De Cara 2009). Specifically, in our focal species, previous work using candidate genes for drought stress provides evidence for divergent selection driving speciation, despite low genome-wide levels of differentiation (Moreno-Letelier & Barraclough 2015). Although a thorough examination of exome-wide variation remains to be done, the correlation of *h* with drought related variables when coupled with the work of Moreno-Letelier & Barraclough (2015) implies that adaptive responses to drought stress likely contributed to the origin and maintenance of species boundaries in this system.

A positive correlation between the steepness of genomic clines (*β*) and *F*_CT_ points towards coincidence of loci involved in disruptive selection and those involved in reproductive isolation. Such a positive association has been demonstrated across several taxa (*cƒ*. Janoušek *et al.* 2012; Parchman *et al.* 2013; Gompert *et al.* 2014; Ryan *et al.* 2017) and we suggest it to be indicative of disruptive selection driving the evolution of intrinsic barriers and its coupling with extrinsic processes. Under the demographic scenario of ongoing gene flow, signatures of selection against hybrids (i.e., underdominance) would be reflected by steep genomic clines (positive *β*), while selection for hybrids (i.e., overdominance) would be reflected by wide genomic clines (negative *β*; Gompert & Buerkle 2011; Janoušek *et al.* 2012). The observed absence of positive *β* outliers and of islands of divergence in our demographic analysis indicates that despite some evidence of coupling between intrinsic and extrinsic barriers, widespread intrinsic incompatibilities are absent in this system, at least for the loci examined in this study. This is consistent with known patterns of forced crosses for these and other white pine species (Critchfield 1986). The limited evidence of intrinsic incompatibilities noted in our study could be generalized across conifers with similar divergence history, owing to their life history strategies such as long generation time and high dispersal capacity, which will restrict the evolution of post- and pre-zygotic isolating mechanism (Stacy *et al.* 2017). Absence of negative *β* outliers and of recent hybrids indicates widespread recombination within the hybrid zone and an intermediate stage of divergence between our focal species (Nosil *et al.* 2009). The intermediate stage of divergence between our focal species, despite a long period of divergence in absolute time (i.e. years), is not surprising given the long generation times and large *N*_e_ estimates for conifers, which would have reduced the realized period of divergence when measured in coalescent units. Overall, the total absence of *β* outliers indicates a viable hybrid zone maintained largely through extrinsic factors (Kamdem *et al.* 2016), which may be the first stage of coupling between intrinsic and extrinsic barriers.

Contrary to the absence of *β* outliers, we identified many *α* outliers which is reflective of a hybrid zone experiencing moderate selection pressure and high levels of gene flow from the parental species (Gompert & Buerkle 2011). Limited variation in *β* is associated with a diffuse genomic architecture of isolation (Gompert *et al.* 2012b), whereas the high genomic heterogeneity in *α*, under the estimated demographic scenario, could imply divergent natural selection operating within the hybrid zone (Gompert & Buerkle 2011). This agrees with the higher values of multilocus *F*_*ST*_ within the putative hybrid zone (*F*_*ST*-periphery_) and previous evidence of local adaptation in this region (Goodrich *et al.* 2016). A similar genomic mosaic of introgression has been noted across several studies (Lexer *et al.* 2010; Parchman *et al.* 2013; Gompert *et al.* 2014; Lindtke *et al.* 2014; de Lafontaine *et al.* 2015) and is likely a result of complex interactions between divergence history, selection, and genomic features.

Evidence of higher number of outliers from *P. strobiformis* and a negative association between our cline parameters (*α* and *β*) could be explained by three processes: (i) intrinsic incompatibilities resulting from Dobzhansky–Muller effects or more complex epistatic effects disproportionally favoring allelic combinations from *P. strobiformis* in the hybrids relative to *P. flexilis* parental background, (ii) widespread directional selection on alleles from *P. strobiformis* in the hybrid zone leading to the formation of co-adapted gene complexes, and (iii) incomplete lineage sorting resulting from recent divergence between Core and Periphery. In contrast to inferences from the Engelmann–white spruce hybrid zone (De La Torre *et al.* 2014b), the asymmetry of outlier loci is not due to high rates of gene flow from Core into Periphery, as the best demographic model excluded gene flow between these groups (see Figure 5B). A higher number of outlier loci with introgression favoring *P. strobiformis* is consistent with the strong influence of selection favoring alleles with *P. strobiformis* ancestry in the hybrid zone. Even without a linkage map, the cline results, along with precipitation seasonality being a strong shared niche predictor for Core and Periphery, points towards widespread directional introgression from *P. strobiformis* into the hybrid zone, which is consistent with local adaptation driving the evolution of co-adapted gene complexes from *P. strobiformis* and of emerging intrinsic incompatibilities (Gompert *et al.* 2012b). The geographic clines of *h*, despite the absence of current gene flow between the Core and Periphery, also points towards an effect of incomplete lineage sorting. However, higher directional introgression from *P. strobiformis* even after accounting for the skewed pattern of genomic ancestry in the hybrid individuals emphasizes the role of selection over incomplete lineage sorting. Further, directional introgression of alleles from Core into Periphery might also have led to the asymmetrical niche divergence between these groups.

Our results are in accordance with studies in other coniferous species demonstrating that speciation is likely initiated through ecological barriers, and several generations of hybridization might occur before the evolution of intrinsic barriers to gene flow (Hamilton *et al.* 2013; Zhou *et al.* 2014; Stacy *et al.* 2017). Integrating the existing genomic dataset with ongoing planting experiments involving climate treatments and measurements of fitness related traits should also help resolve the joint influence of extrinsic and intrinsic isolating mechanisms. Specifically, coincidence between the steepness of genomic, geographic, and trait specific clines would indicate a dominant role of extrinsic factors in facilitating divergence and speciation (Holliday *et al.* 2010; De La Torre *et al.* 2015; Stankowski *et al.* 2015; Ryan *et al.* 2017). Alternatively, the presence of several loci showing steep clines, but lacking climatic or functional associations would indicate a dominance of intrinsic barriers (Ryan *et al.* 2017). Although the genomic cline analysis used in this study provided key insights into a complexity of species isolation, it lacks sufficient power to account for complex epistatic effects (Gompert & Buerkle 2011). These have likely played a key role in ecological speciation and in initiating the evolution of reproductive isolation (Lindtke *et al.* 2012; Flaxman *et al.* 2014). Ultimately, furthering genomic resources will help test whether absolute measures of divergence are correlated with recombination rate, and will account for the non-random genomic distribution of climate-associated genes and their tendency to co-localize in areas of reduced recombination (Wolf & Ellegren 2016). This study, however, provides concrete evidence of ecological speciation with gene flow, the presence of a historical hybrid zone maintained by extrinsic factors, and early stages of coupling between disruptive selection and intrinsic barriers contributing towards diversification. Whether these patterns hold generally for speciation within conifers, given their life history characteristics as well as their complex and large genomes, is thus a worthwhile area of future research.

## Acknowledgements

This research was funded by U.S. National Science Foundation (NSF) grants EF-1442486 (Eckert), and EF-1442597 (Waring), as well as by Northern Arizona University’s Technology Research Initiative Program, USDA Forest Service Forest Health Protection Gene conservation program, USDA Forest Service National Fire Plan Award 01.RMRS.B.6 (Schoettle), Virginia Commonwealth University (VCU) Department of Biology, and VCU Integrated Life Sciences. We thank Z. Gompert and T. Parchman for providing R code that aided plotting the BGC results. We also acknowledge our field sampling and lab work crew (see Supporting information, Appendix S1.D). This work would not have been possible without generous computational resources provided by the VCU Center for High Performance Computing and the Brigham Young University Fulton Supercomputing Lab.

## Data accessibility

Raw reads generated during this study are available at NCBI SRA database (SRXXXXXXX). Genotype file following SNP calling in 012 format and admixture proportions per individual tree are available at Dryad (doi: XXXX).

## Author contributions

The study was designed by KMW, AVW, AJE, LFR, CW, and SC. Field sampling was performed by AWS, FMF, LFR, MSG, CW, ALS, and KMW. Funding for this study was procured by KMW, AWS, and AJE. MM, JCB and CF performed the data analysis. MM generated the genomic data and wrote the manuscript. All authors edited the article and have approved the version for submission.

## Supporting Information

**Table S1.** Sampling locations used in this study, their classification into Core and Periphery, and mean admixture proportions.

**Table S2**. Raw and converted parameter estimates from the ∂A∂I model that was best-supported by AIC model selection.

**Fig. S1.** Schematics and parameter details for each of the 11 demographic models for the divergence of core and periphery groups within *P. strobiformis* and *P. flexilis* run in our ∂A∂I analysis. Parameters include divergence times (*T*_i_), population sizes (*N*_i_), homogeneous rates of gene flow (*M*_ij_, gene flow from lineage j to i) and genomically heterogeneous rates of gene flow (*M*_ijh_).

**Fig. S2.** Ecological niche model projections for Core, Periphery, and *P. flexilis*, under present and past climate

**Fig. S3. A**) Climate PCA with variables used in the ENMs **B)** Distribution of precipitation seasonality (Bio15) at presence locations of Core & Periphery.

**Appendix S1.A)** Filtering of occurrence records and ENM details, **B)** Bioinformatic pipeline for processing raw ddRADseq reads, and **C)** Down-sampling of SNP loci for demographic modeling using ∂A∂I. **D)** List of volunteers and technicians involved in sampling and library preparation.

---

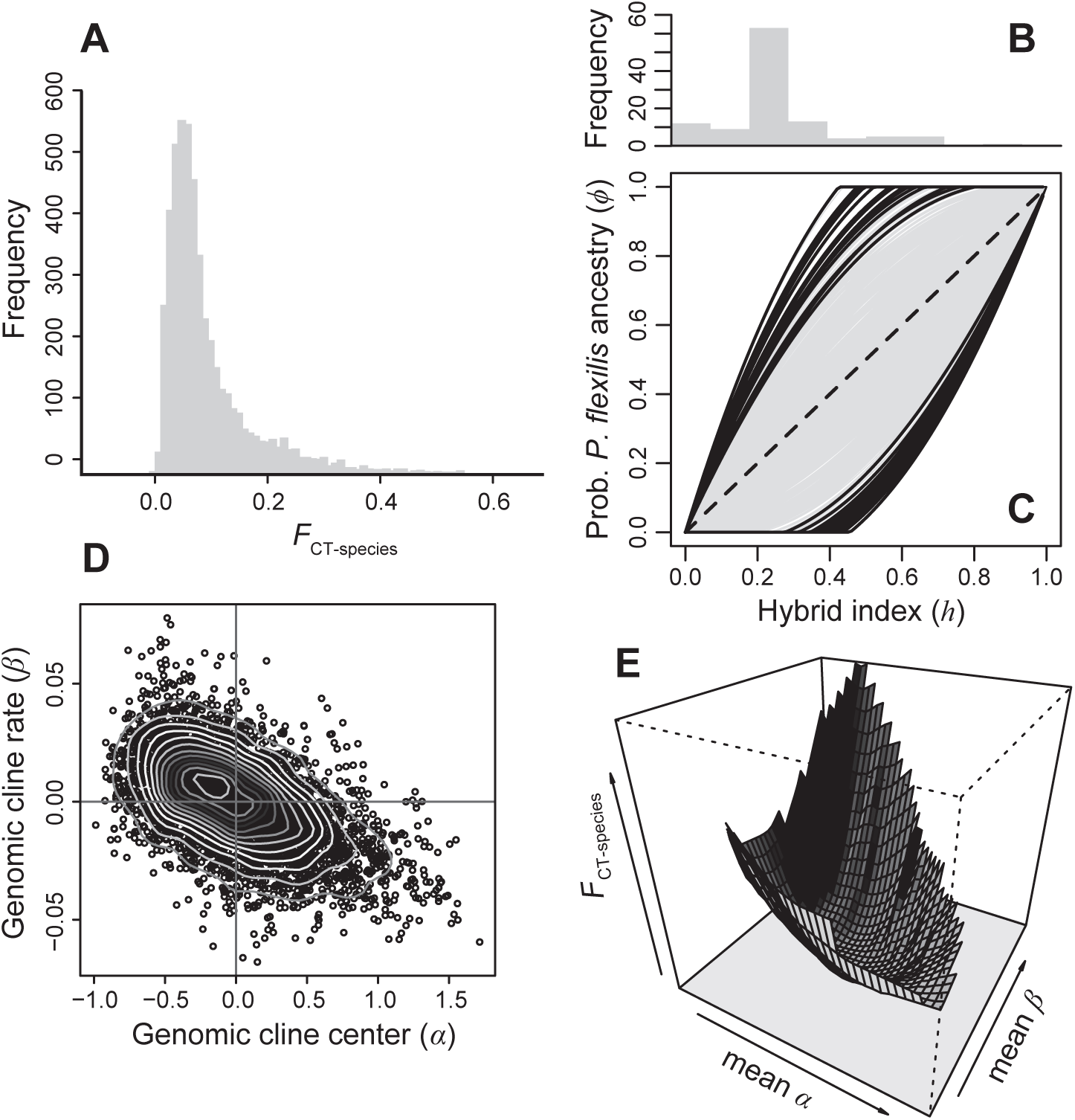

